# In Vitro and In Vivo Pathogenicity of *A. alternata* to Sugar beet and Assessment of Sensitivity to Fungicides

**DOI:** 10.1101/2022.01.15.476474

**Authors:** Md Ehsanul Haque, Most Shanaj Parvin

## Abstract

*A. alternata* is a weak and opportunistic pathogen, but recently this pathogen has been reported in major sugar beet growing regions in the US. Therefore, we were interested to investigate the in-vitro and in-vivo pathogenesis of *A. alternata* to sugar beet seed, seedlings, tissue slices, stecklings, and roots. The in-vitro study demonstrated that *A. alternata* reduces seed germination, causes damping-off of seedlings, and decomposes beet tissue. The in-vivo study showed the same inoculum responsible for Alternaria leaf spots, and black circular spots on beetroot. Furthermore, we did a preliminary study to determine in vitro sensitivities of some fungicides to determine the effective concentration that inhibited 50% of radial mycelial growth (EC50 values) of *A. alternata*. The results indicated that EC50 values ranged from 0.0023 to 0.0058 for PROLINE, from 0.0004 to 0.0021 for OMEGA, and from 0.0027 to 0.0053 for Super TIN. Among the potential fungicides, the efficiency of quinone outside inhibitors (QoI- PRIAXOR) was relatively higher to inhibit the radial growth of *A. alternata*, irrespective of concentrations.

## 1. Introduction

*A. alternata* (Fr.) Keissler (anamorph) is regarded as an important saprophyte, but also a necrotrophic plant pathogen on multiple hosts over 380 including sugar beet [19,18]. It can be found in plants, soil, food, and indoor air in temperate and tropical regions. It causes leaf spots, rot, and blight in many plant parts. Macroscopically, it is a fast-growing colony, whitish at the border and light greenish to olivaceous brown at the young stage, grey to black, and powdery to floccose at maturity. Conidiophore septate and hold chain of conidia (4-8 conidia), conidia obclavate shaped with a short beak and contains several transverse and longitudinal septa [20, 21, 22, 4].

*A. alternata* favors a moist warm environment for sporulation and a cool and humid environment for new infection. Conidia and conidiophores carry over in plant debris, seed, and seedlings and can spread through wind and rain splash. *A. alternata* infects plants through wounds, lenticels, stem ends, pedicels, and cuticle using its appressorium [14, 10, 9]. It also can cause disease when mechanical infection happens due to virus infection [17], drought damage, and lack of nutrients in the host plant.

An Alternaria Leaf Spot of the Sugar Beet caused by *A. brassicae* was first reported in California [1]. It was of little importance in the sugar beet crop in the US but nowadays, it has drawn the attention of the growers and researchers due to the increased incidence and severity of this pathogen since 2010. Recently, *A. tenuissima* and *A. alternata* were reported to cause leaf spots on sugar beet in Minnesota and North Dakota, respectively [4, 23]. The most common method for controlling Alternaria diseases is the use of fungicides both in fields and in storages of various economic plants [15, 13, 16, 2]. The development of resistance in pathogenic fungi to common fungicides and increasing residual hazardous effects on human health and environmental pollution has given a thrust to search for new derivatives that can obstruct the fungal pathogenicity [12]. Fungicide-resistant to *A. alternata* impedes the practical control of the Alternaria diseases in crop fields [3]. The effectiveness of several groups of fungicides varied significantly. Therefore, the present work aimed to study the in- vitro assay of some selected fungicides against *A. alternata*.

## 2. Materials and Methods

### 2.1. Preparation of Spore Suspension

Clones of *A. alternata* (Genbank accession: MK439485.1), was maintained on 50% Potato Dextrose Agar (PDA) (https://www.ncbi.nlm.nih.gov/nuccore/MK439485.1). The colony showed grey to olive green morphology and microscopically it indicated a long chain of conidia. A single spore (conidium) of *A. alternata* was isolated and sub-cultured. Genomic DNA was extracted from these pure cultures, and molecular assay using ITS 4 and ITS5 confirmed these cultures as *A. alternata* which were used for in vitro and in vivo study.

Sterile distilled water was added to plates containing 10 days old *A. alternata* culture and then the surface of the agar was scraped with a sterilized spatula to remove hyphae and spores. The contents of several plates were poured through cheesecloth into a beaker to remove agar and large fragments of mycelium. The spore concentration was determined with a hemocytometer and adjusted to approximately 5 × 10^5^ spores per ml of *A. alternata*.

### 2.2. Inoculation of Sugar beet Seed with A. alternata

Fungicidal coats were removed from the seed, surface sterilized with 10% bleach solution with a drop of Tween-20, washed in autoclaved water, removed excess moisture, and dried inside the laminar airflow cabinet. Fifty seeds were treated or soaked overnight with a spore suspension of *A. alternata* (5 × 10^5^ spores per ml/Petri dish) and kept at 24° C and 70-80% relative humidity. Mock treatment was performed with autoclaved water with the same number of seeds. Germination records were collected 7 days after post-inoculation.

Alternatively, to check seed germination under *A. alternata* inoculation seeds were washed and surface sterilized with a bleach solution: 10%, subsequently, blotter paper technique was applied following International Seed Testing Association (ISTA) guideline. Autoclaved blotter paper was, folded four times, each fold contained 25 seeds and inoculated with *A. alternata* spore suspension. A hundred seeds for both non-inoculated and inoculated were taken for quantifying the germination percentage. Data were recorded at 7 days after inoculation.

### 2.3. In-vitro Inoculation of A. alternata on Seedlings Grown in Falcon Tube

Surface sterilized seeds were gown in 50 ml falcon tube containing 50% PDA media, and allowed to grow for 5 days. After 5 days of seedlings’ age, each seedling was inoculated with *A. alternata* spore suspension (5 × 10^5^ spores per ml) @ 100 µl/plant except the control plant. Data were recorded nine days after inoculation.

### 2.4. In-vitro Inoculation of A. alternata on Fresh Healthy Tissue

Fresh healthy tissue of sugar beet was sliced into pieces, surface sterilized with 10% bleach solution, washed in autoclaved water, removed excess moisture with autoclaved Whatman paper. Spore suspension (5 × 10^5^ spores per ml) conidia of *A. alternata* was inoculated onto beet tissues. Data was recorded three times such as 7, 10, and 15 days after inoculation. It indicated considerable changes in beet tissue color and texture at different days after inoculation due to infection and resulted in decomposition and discoloration of tissue except for the control plate.

### 2.5. Greenhouse Evaluation of A. alternata to Sugar beet

*A. alternata* spore suspension (5 × 10^5^ spores per ml) was sprayed twice to sugar beet leaves and root zone first at four weeks and second at six weeks of plant age at 25° C and 80% relative humidity. Eight plants were inoculated except control plants. Data were recorded at 16 weeks of post-inoculation.

### 2.6. Fungal Growth Inhibition Study via Fungicides

Preparation of media: 15 g Agar powder, 12 g potato dextrose broth, and 1000ml water used for preparing 50% potato dextrose agar (PDA) media, autoclaved and amended with antibiotics, and the desired concentration of fungicides included 1 ppm, 0.1 ppm, 0.01 ppm, 0.001 ppm, and 0 ppm as control. Media was poured in a 90 mm Petri dish with three replications for each concentration and allowed to solidify (Table 1). The relative efficacy of four (04) different fungicides was tested against *A. alternata* in the laboratory. Fungal plugs were cut out from the fungal culture using a 5-mm- diameter corker borer and placed on 50% PDA supplemented Petri dishes with 1ppm, 0.1ppm, 0.01ppm, 0.001ppm in triplicates. *A. alternata* inoculum at least 8-10 days old culture was used and placed in the center of each Petri dish. The medium without any fungicide poured and inoculated similarly served as control. The Petri dishes were incubated at 28±1°C for 12 days. The efficiency of fungicides was assessed by measuring the radial growth of the fungal colony in cm.

**Table 1.** Fungicides tested for in-vitro evaluation against *A. alternate*

| Fungicides | Active ingredient | Rate/Dose |
| --- | --- | --- |
| Super Tin | Triphenyl Tin Hydroxide | 1ppm, 0.1ppm,<br>0.01ppm, 0.001ppm |
| Omega | Fluazinam |  |
| Priaxor | Fluxapyroxad and<br>pyraclostrobin |  |
| Proline | Prothioconazole |  |

#### Statistical analysis

Statistical analysis consisted of one-way ANOVA with Tukey’s or Sidak’s multiple comparisons testing performed by GraphPad PRISM 9.3.1. All experiments were done in triplicates and the data is plotted as the mean ±SE of triplicate determinations. EC50 calculation was also performed by GraphPad PRISM 9.3.1 via normalization and nonlinear regression curve fit.

## 3. Results

### 3.1. In vitro Inoculation- A. alternata Reduced Sugar beet Seed Germination

In vitro inoculation via conidia suspension of sugar beet seeds demonstrated below 50% germination, and sprouted seeds were rotten. In mock inoculation, above 90% of seeds were sprouted and normal growth was observed (Fig. 1 A & B). Likewise, blotter paper technique in non-inoculated check-showed 90% germination while paper contained seed and conidial suspension of *A. altarnata* showed only 20% germination (Fig. 2 C & D).

**Fig 1.**
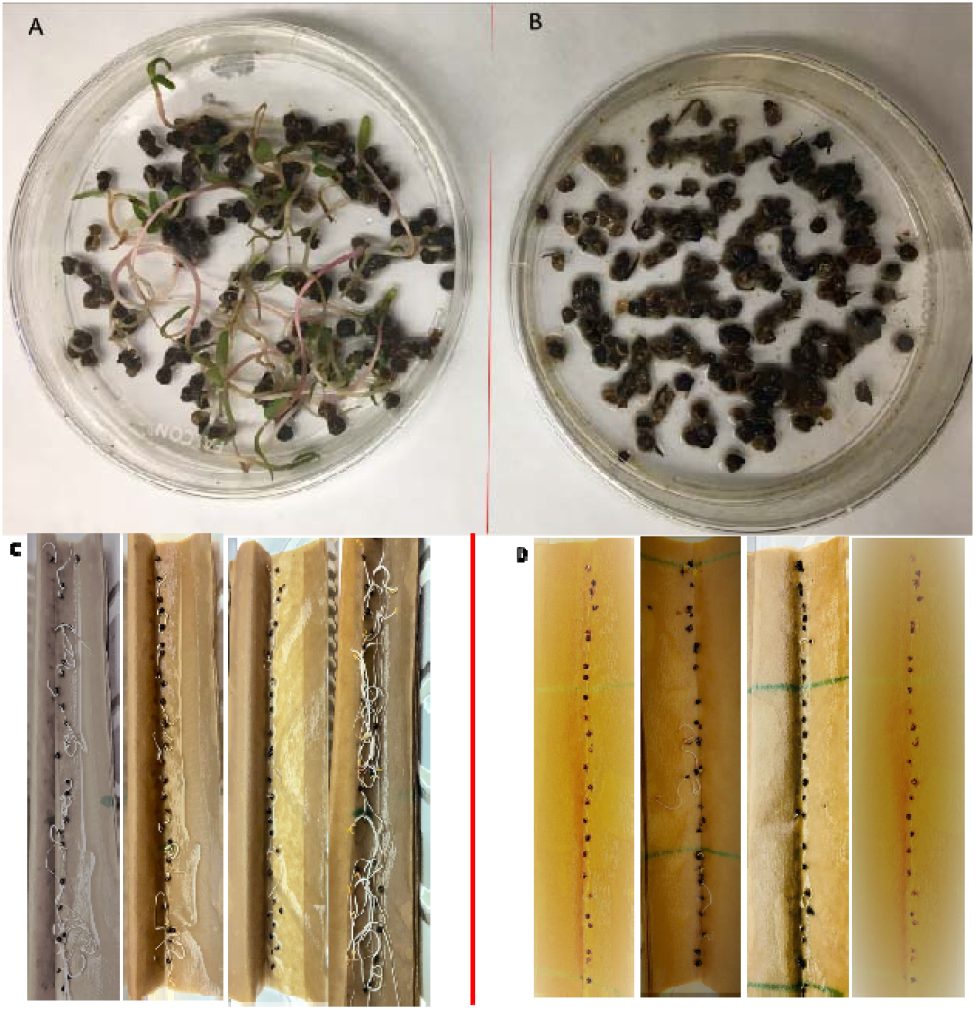
In vitro conidial-inoculation and mock-inoculation of sugar beet seed on Petri plate and Blotter paper at 7 dpi- A. plate contained only seed (non-inoculated check)- shows 90% germination, B. plate contained seed and conidial suspension of *A. altarnata*- shows below 50% germination, C. paper contained only seed (non-inoculated check)-shows 90% germination, D. paper contained seed and conidial suspension of *A. altarnata* -shows 20% germination.

**Fig 2.**
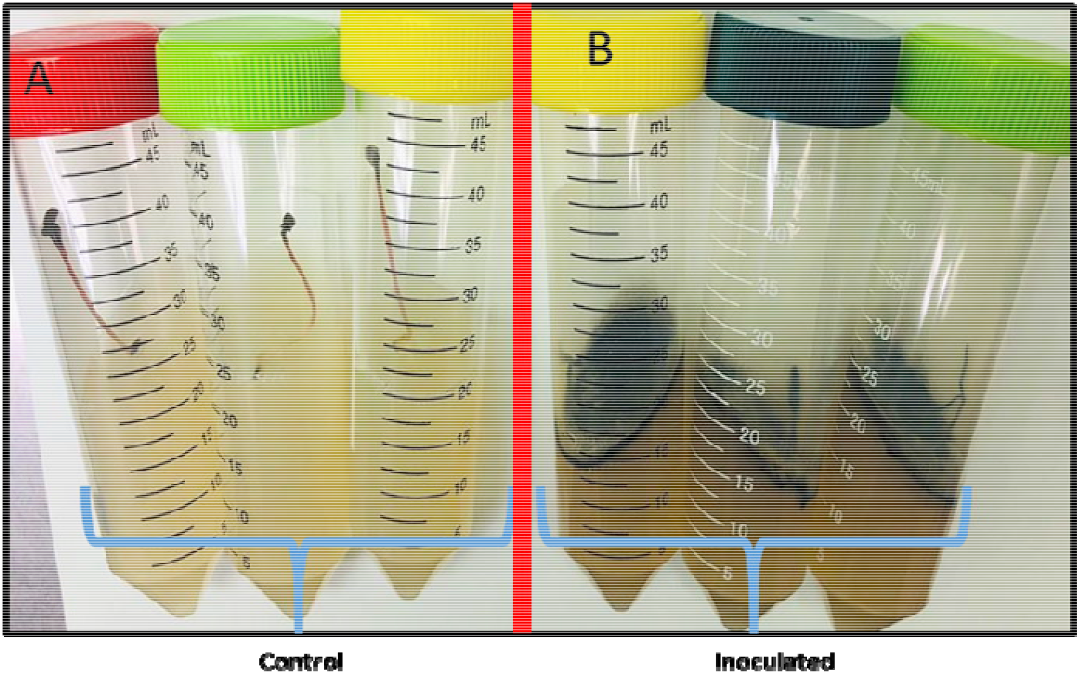
In-vitro inoculation of *A. alternata* in seedlings grown in falcon tube A. non- inoculated check-sprouted seedling shows normal growth, B. seedling death occurred in *A. alternata* inoculated tubes at 9 days after post-inoculation.

### 3.2. In-vitro Inoculation of A. alternata in Seedlings Demonstrated Damping-off Like Symptoms

In order to understand the pathogenesis of *A. alternate*, the seedling was inoculated ambient condition in a falcon tube. Nine days after inoculation all plants were infected and died off except non-inoculated control (Fig. 2).

### 3.3. In-Vitro Inoculation of A. alternata on Fresh Sugar beet Tissue

In order to understand the pathogenesis of *A. alternate*, sugar beet slices were inoculated at the inoculation chamber. This shows infection like tissue rotten in inoculated sugar beet slices. The infection resulted in the discoloration of tissue which is followed by decomposition (Fig. 3 B). There was no tissue decomposition in the control plate (Fig. 3 A).

**Fig 3.**
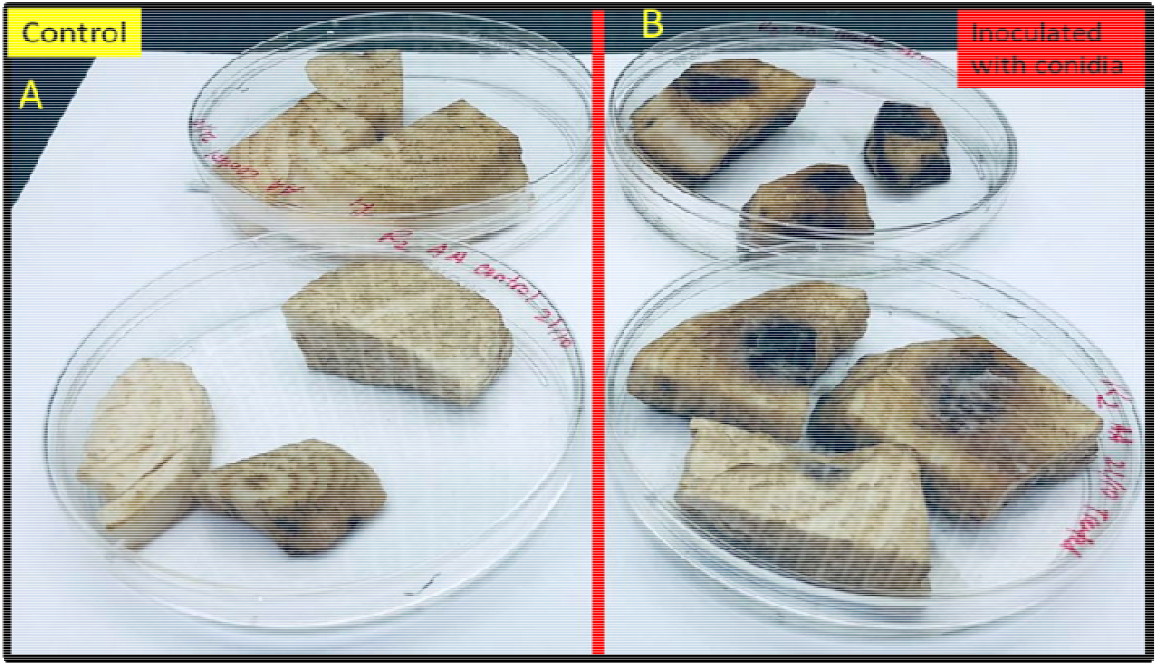
In-vitro inoculation of *A. alternata* on sugar beet slices A. non-inoculated check- sugar beet slice shows zero rotten, B. inoculated slices shows black dark rotten signs at 3 weeks after post-inoculation.

### 3.4. In Vivo Study Showed Pathogenicity of A. alternata

A greenhouse study was performed to understand the pathogenesis of Alternaria to sugar beet plants. 16 weeks after post-inoculation, characteristics circular brown lesions appeared on the older leaves (Fig. 4 A & B), while the non-inoculated plants did not show any symptoms. Subsequently, sugar beet roots were harvested and the dark brown circular symptoms found on the surface of beet appeared on 30% of the harvested beet (Fig. 4 D). No symptoms were observed on control beets. The experiment was conducted twice.

**Fig 4.**
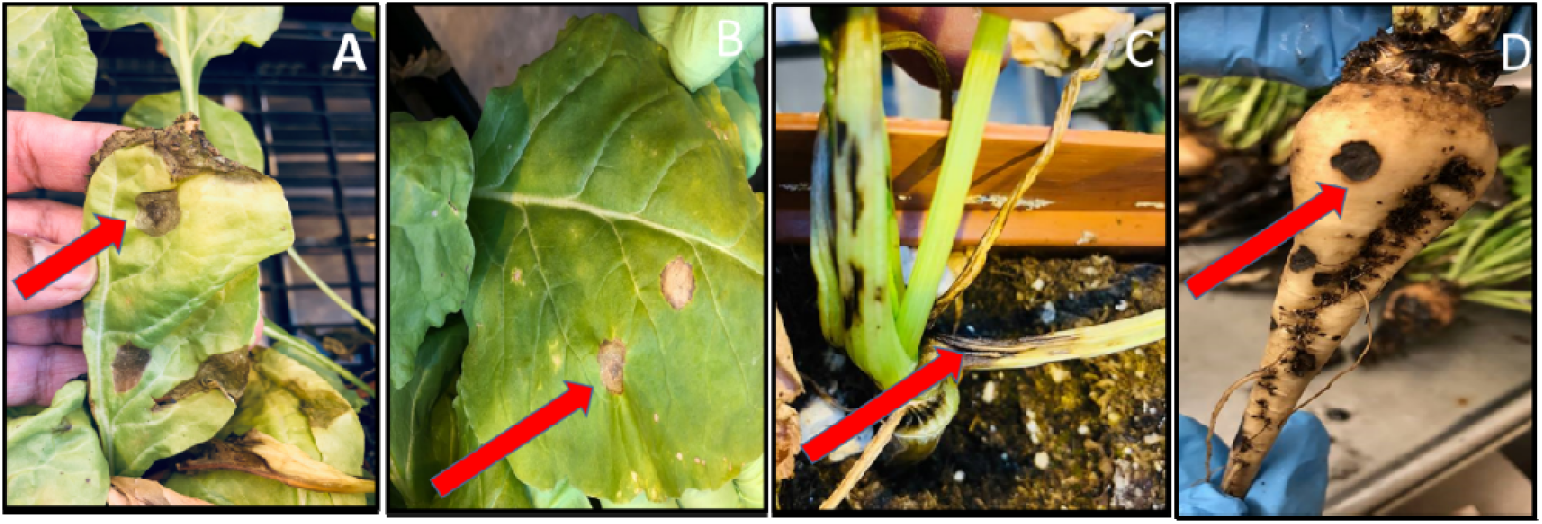
*A. alternata* symptoms appeared on leaf, stem, and sugar beet root after 16 weeks of post-inoculation in the greenhouse. The red arrow indicates typical signs of infection.

**Fig 5.**
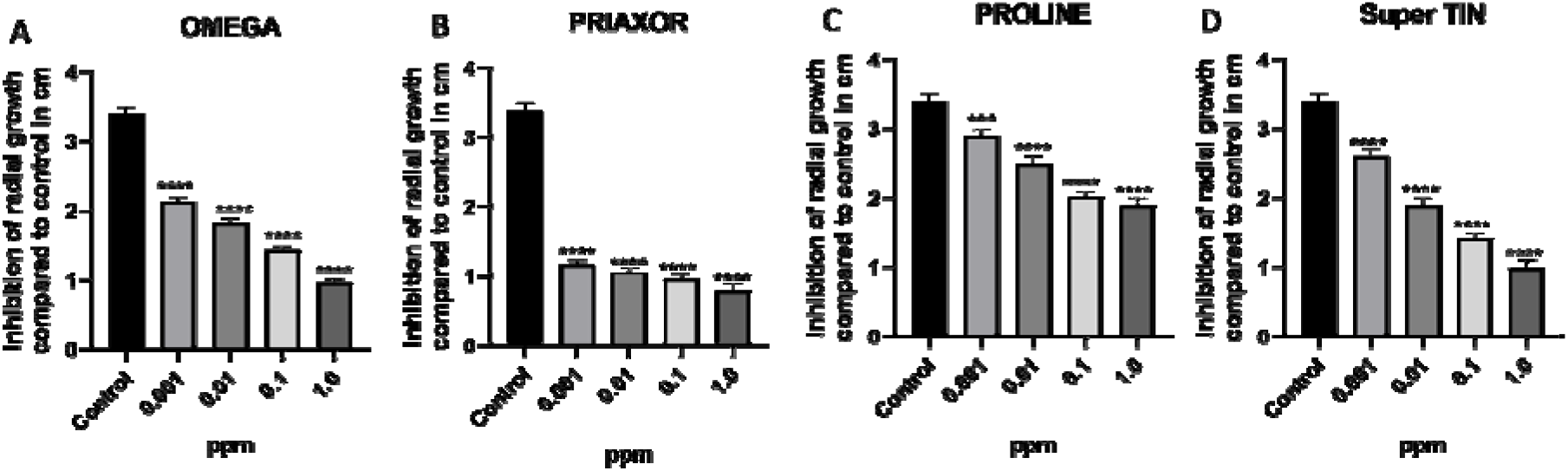
Radial growth (cm) of *A. alternata* compared to non-treated mock with four fungicides with four concentrations at 0.001 ppm, 0.01 ppm, 0.1 ppm, and 1.0 ppm. Radial growth of (A) OMEGA, (B) PRIAXOR, (C) PROLINE, and (D) Super TIN. ****; ***; **; * indicates significant differences in radial growth compared to the control 0 ppm concentration at p- value of ≤ 0.0001; ≤ 0.001; ≤ 0.01; ≤ 0.05 respectively.

### 3.5. Radial Growth and EC50

In vitro efficacy of quinone outside inhibitors (QoI- PRIAXOR) was relatively higher to inhibit the radial growth of *A. alternata* compared to the other three fungicides, irrespective of concentrations (Fig. 4 B). Triphenyl Tin Hydroxide (Super TIN) and Omega showed moderate efficiency in controlling the radial growth (Fig. 4 A & D). Triazole (PROLINE) showed minimum efficiency while compared to the non-treated check (Fig. 4 C).

EC50 (the effective concentrations to cause inhibitions by 50) were used to understand the fungicide potency. Among the four fungicides, three fungicides showed the sigmoid curve or S-shaped curve. This shows a growth pattern of A. alternate where the radial growth slowly reduced with the increasing concentration of fungicides on the culture plates. Super TIN EC 50 was 0.00385, with an R squared value 0.97, which is followed by PROLINE (EC50 = 0.003798), with an R squared value 0.95. The Goodness of Fit was very close to 1 for Super TIN and PROLINE. The EC50 of OMEGA was 0.0011, with an R squared value 0.91 (Fig. 6).

**Fig 6.**
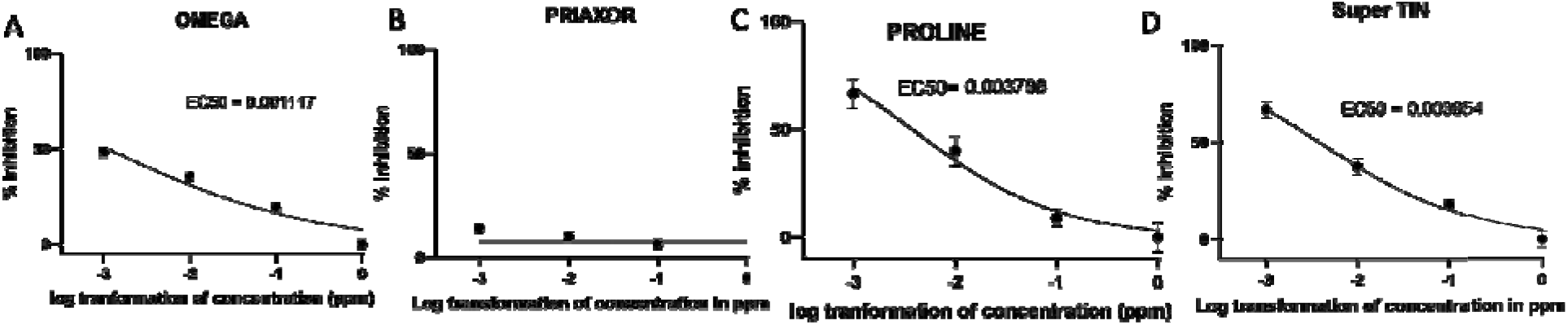
EC50 values for four fungicides and log transformation value four concentrations at 0.001 ppm (−3), 0.01 ppm (−2), 0.1 ppm (−1), and 1.0 ppm (0) in *A. alternata*. EC50 of (A) OMEGA = 0.001117, (B) PRIAXOR = infinity, (C) PROLINE = 0.003798, and (D) Super TIN = 0.003854.

## 4. Discussion

This study provided in vitro evidence of *A. alternata* reduced sugar beet seed germination and sprouting. In situ is relatively heterogeneous and often does reflect the true causal organism that inhibits the seed germination. Our study demonstrated that *A. alternata* can decrease sugar beet seed germination by 50%.

*In-vitro* inoculation of *A. alternata* in seedlings showed that *A. alternata* is involved in damping-off-like symptoms in sugar beet. Previously we learned that damping-off symptoms are mostly caused by *Aphanomyces cochlioides, Rhizoctonia solani*, and *Globisporangium ultimum* [5,6,7]. Our study found that susceptible sugar beet cultivar might incline to damping-off disease if the field sustains a previous story of *A. alternata*.

In-vitro inoculation of *A. alternata* on fresh sugar beet tissue demonstrated that *A. altarnata* can be an emerging post-harvest pathogen. Sugar beet piled for a long period of time before processing for sugar. Thus, it might be an issue with the storage of sugar beet. In this in-vitro pathosystem, we noticed the potential of *A. alternata* to cause tissue discoloration and decomposition of sugar beet. Recently, several new post-harvest pathogens have been reported in sugar beet such as *Talaromyces pinophilus, Geotrichum candidum*, and *Clonostachys rosea* [8, 24, 25]. Our study shows that Alternaria decomposes sugar beet tissue, which can be a potential threat for storage beets.

Previously it was known that Alternaria causes Alternaria Leaf Spot of the Sugar Beet. Our in vivo study showed the pathogenesis of *A. alternata* causes not only leaf spots but also causes brown lesions on the collar region and dark brown circular spots on stecklings.

Emerging pathogen is always a concern for growers because of fungicide sensitivity issues and fungal resistance via mutation to a particular group of fungicides. Moreover, biological control often does not work managing new pathogens in field conditions. Therefore, there is no feasible alternative management approach apart from chemical fungicides. Our preliminary study on Alternaria-fungicide combination via radial growth measurement and EC50 values estimation showed that Priaxor, Super Tin fungicides can be recommended to effectively control this pathogen.

## 5. Conclusion

*A. alternata* becoming prevalent along with Cercospora leaf spot. *Alternaria* species were reported from sugar beet in California, Michigan, Montana, North Dakota, Arizona, and Minnesota. It might be a problem for the storage of sugar beet. Careful monitoring, clean cultivation, and seed treatment are imperative for handling the disease. Priaxor, Omega, and Super Tin fungicides can be recommended to effectively control this pathogen.

## Acknowledgment

Sugar beet Research and Education Board and American Sugar beet Growers Association.

